# Deciphering functional tumor-immune crosstalk through highly multiplexed imaging and deep visual proteomics

**DOI:** 10.1101/2024.05.22.595266

**Authors:** Xiang Zheng, Andreas Mund, Matthias Mann

**Affiliations:** 1: Novo Nordisk Foundation Center for Protein Research, University of Copenhagen, Copenhagen, Denmark; 2: Proteomics and Signal Transduction, Max Planck Institute of Biochemistry, Martinsried, Germany

**Keywords:** Multiplex staining, mass spectrometry, deep visual proteomics, tumor microenvironment, cancer

## Abstract

**Deciphering the intricate tumor-immune interactions within the microenvironment is crucial for advancing cancer immunotherapy. Here, we developed a novel approach integrating highly multiplexed imaging, laser microdissection, and deep visual proteomics (DVP) to spatially profile the proteomes of 21 distinct cell populations in human colorectal and tonsil cancers with high sensitivity. We selected colorectal tumor as an example of a cold tumor, uncovering an immunosuppressive macrophage barrier impeding T cell infiltration and regionally altering lymphocyte proteomes. Spatial proteomic analysis revealed distinct functional states of T cells in different tumor compartments. In tonsil cancer - a hot tumor, we identified significant heterogeneity of cell proteomes influenced by proximity to cytotoxic T cell subtypes. Tumor-infiltrating T cells exhibited metabolic adaptations and enhanced stress resilience, enabling migration into hypoxic regions. Our spatially-resolved, highly multiplexed strategy deciphers the complex cellular interplay within the tumor microenvironment, with implications for identifying new immunotherapy targets and signatures.**

**Highlights:**

- Novel approach integrates multiplexed imaging and deep proteomics to profile the tumor microenvironment
- Identified 21 distinct cell populations and thousands of proteins from single cell types
- Macrophage barrier impedes T cell tumor infiltration and T cells adapt to hypoxia
- Spatial insights may guide new immunotherapy targets and personalized oncology

## Introduction

The tumor microenvironment (TME) is a complex ecosystem where cancer cells dynamically interact with diverse non-malignant cells, including immune cells, fibroblasts, and endothelial cells, as well as the extracellular matrix ^1^. These intricate interactions shape all stages of cancer progression, from tumor initiation and growth to metastatic spread, and profoundly influence responses to therapy ^2^. In particular, cross-talk between cancer cells and immune cells within the TME plays a central role in modulating anti-tumor immunity, often leading to immunosuppression that enables tumor immune evasion ^3^. Deciphering the cellular composition, spatial organization, and molecular circuitry of the TME is therefore crucial for understanding tumor biology and developing effective immunotherapies.

Advances in single-cell technologies have revolutionized our ability to dissect the cellular heterogeneity of tumors ^4,5^. Single-cell RNA sequencing (scRNA-seq) enables unbiased transcriptional profiling of individual cells within complex mixtures, revealing rare and novel cell types and states. However, scRNA-seq requires tissue dissociation, which leads to loss of spatial information ^6^. Spatially resolved transcriptomics methods like MERFISH and seqFISH have emerged to map single-cell gene expression in situ, but they currently profile a limited number of pre-selected genes ^7^. At the protein level, while imaging-based approaches like multiplexed ion beam imaging (MIBI) and imaging mass cytometry (IMC) can spatially profile targeted proteins, their limited depth hinders unbiased discovery of the intricate cellular networks within the TME ^8^. Conventional single or low-plex staining methods often fall short in retaining spatial relationships across multiple cell populations and they do not achieve the highly multiplexed characterization needed to capture the full complexity of the TME ^8,9^. Spatial omics has been combined with antibody-based read outs which targeted to a limited number of antigens, which may not have been validated in different tissues and not yet in a deep and unbiased proteomics format^4^

Mass spectrometry (MS)-based proteomics has long been used to characterize tumor tissue at the bulk level with notable successes and with the Clinical Proteomic Tumor Analysis Consortium (CPTAC) as a prominent example ^10–12^. For example, CPTAC recently released a pan-cancer proteogenomic dataset, enabling multi-omics analysis of bulk tumor samples^10,13^. Nevertheless, despite tremendous technological improvement in MS-based proteomics ^14^, all these datasets lack the spatial resolution needed to dissect the complex cellular interplay within the heterogeneous TME at the single cell type level. Achieving highly multiplexed characterization of cells together with deep single cell type proteome characterization requires innovative analytical strategies. We previously introduced deep visual proteomics (DVP), a spatial technology integrating imaging, cell segmentation and classification, laser microdissection, and MS to investigate complex tissues ^15^. However, DVP has only been demonstrated with up to four-plex staining so far.

Here, we aim to further enhance the throughput and depth of DVP by combining it with highly multiplexed imaging as well as multiplexing at the mass spectrometric level in the form of a recently developed multiplexed data independent acquisition (DIA) workflow ^16,17^. By optimizing multiplex staining to maintain morphological integrity, our workflow enables precise laser microdissection and deep spatial proteomics of lymphocytes, overcoming challenges posed by their small size to provide new insights into their complex biology in the native tissue microenvironment ^18^. We evaluated our method’s versatility by applying it to both immunologically cold and hot tumors - those less and more likely to respond to immunotherapy, respectively ^19^ - demonstrating its wide applicability across diverse TME. By applying multiplex staining-powered DVP, we aim to decipher the complex cellular interplay and functional adaptations within the TME, potentially discovering new immunotherapy targets. More broadly, our framework should provide a powerful means for ultra-high sensitivity profiling of limited clinical samples to inform personalized oncology.

## Results

### Integrated 14 to 22-plex imaging and MS for single cell type tumor spatial proteomes

To characterize the spatial molecular landscape of the TME at the single cell level, we first performed 14 to 22-plex cyclic immunofluorescence staining on formalin-fixed paraffin-embedded (FFPE) sections from human colorectal cancer (CRC) and tonsil cancer samples using the MACSima imaging platform (**Figure 1A-C; Figure S1A** and see below). For 22-plex staining this took less than 50 hours on a tissue region of 30 mm^2^, sufficient to isolate even rare cell types from the same tissue slide for downstream analysis.

**Figure 1.**
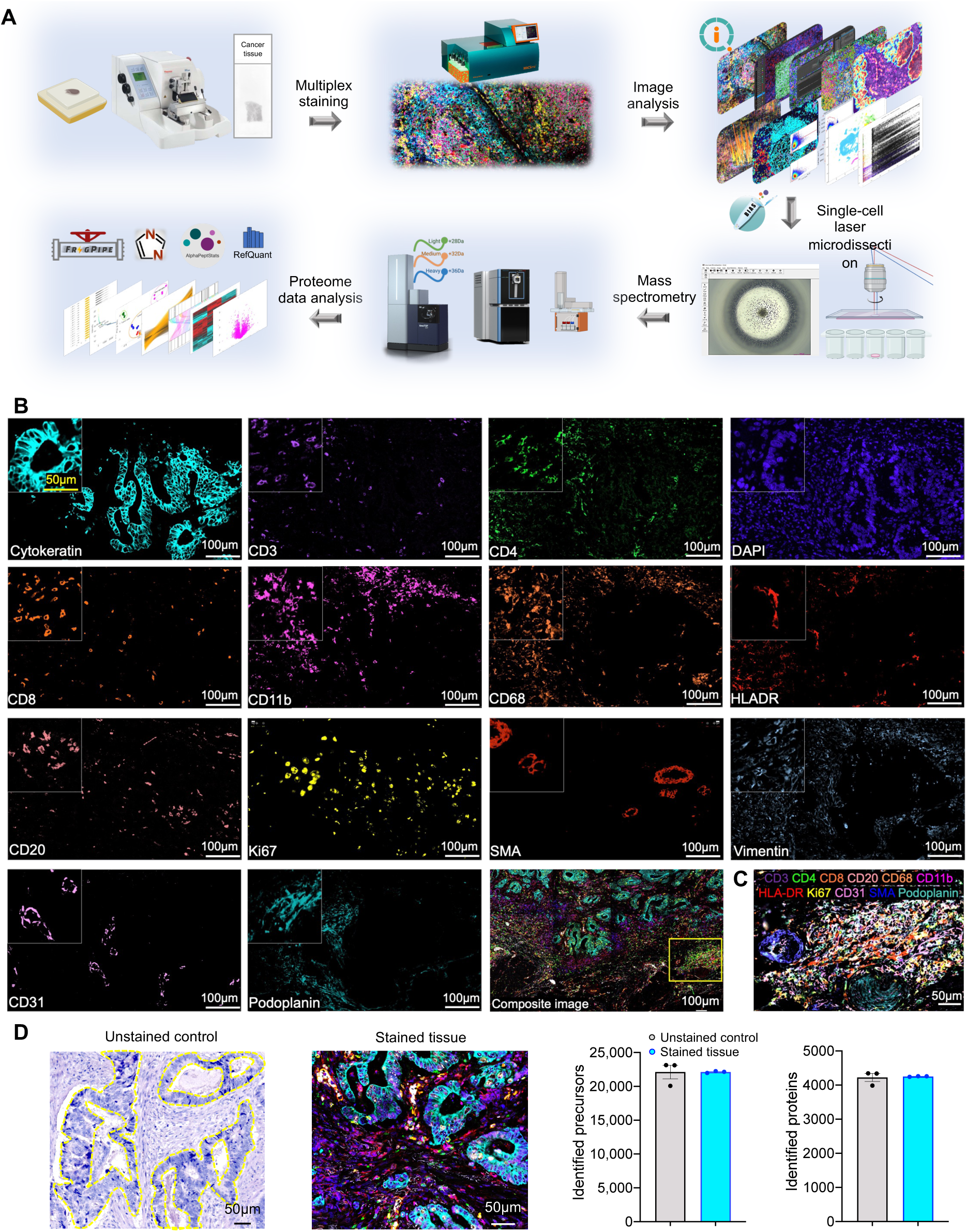
Integrated 14 to 22-plex imaging and MS for single cell type tumor spatial proteomes. *(A) Schematic of the experimental workflow integrating multiplex staining and MS-based proteomics to enable deep proteomic profiling while preserving spatial context of the tissue microenvironment. (B) Representative images: multiplexed pseudo-colored images illustrating marker staining of colorectal cancer FFPE sections. The composite image displays all merged marker staining, providing a comprehensive view of the cellular composition and spatial organization. (C) Representative image of the tertiary lymphoid structure (TLS) outlined in yellow in (B), showing the spatial distribution of immune cells (CD4+ helper T cells, CD8+ cytotoxic T cells, CD20+ B cells, and CD68+ macrophages) and vasculature (CD31+ blood vessels and Podoplanin+ lymphatic vessels). (D) Comparison of the number of precursor and protein identifications of tumor epithelial regions from immunofluorescence-unstained and MACSima-stained tissues, demonstrating minimal protein loss during the staining process. The yellow dotted line indicates the boundary of the tumor epithelial region, while the pseudocolor cyan represents CK+ tumor cells.*

Unlike conventional low-plex staining, multiplex imaging revealed a plethora of multiple cell types and their spatial interactions within the same tissue section (**Figure 1C**). Multiplex staining of CRC tissue facilitated detailed evaluation of the spatial distribution of tumor and stromal cells. Tertiary lymphoid structures (TLS) are important sites of local immune responses within the TME ^20^. Within the lamina propria — a layer beneath the tumor epithelium — we discerned TLS composed of CD4+ helper T cells (TH), CD8+ cytotoxic T cells (CTLs), CD20+ B cells, and CD68+ macrophages. This region also contained CD31+ blood vessels and Podoplanin+ lymphatic vessels. Overall, the tumor epithelium region exhibited fewer immune cell markers compared to the lamina propria, including markers of macrophages, CTLs, TH, B cells, and granulocytes, highlighting spatial heterogeneity in immune cell infiltration.

Next, we utilized high-plex imaging data to perform cell segmentation and classification using the MACS iQ View software ^21^, which was essential for the subsequent laser microdissection and MS-based proteomics analysis (**Experimental Methods**). After preprocessing to stitch images and subtract background noise, we segmented cells by identifying nuclear and specific cytoplasmic markers, with optimizing parameters such as minimum and maximum diameter for cell size, detection sensitivity for maker intensity, separation force for distinguishing adjacent cells, and smoothing filter sigma for reducing noise. Subsequent to segmentation, cells were classified by identifying and categorizing them into distinct types based on their phenotypic markers. We then imported the final classification masks into the BIAS software ^15^ to set up reference points for subsequent automated laser microdissection on a laser microdissection instrument (Leica LMD 7), which took about three seconds per cell shape.

Cell shapes of the same type and region were collected in 384-wells followed by liquid chromatography-mass spectrometry (LC-MS) analysis using very low flow Whisper gradients^22^. For ultimate sensitivity we used both diaPASEF on the timsTOF and the Astral platforms^23,24^. All MS data was analyzed using DIA-NN ^25^. Our workflow analyzed 40 samples per day using a label-free MS method and 80 samples with the dimethyl-labeling method ^16^, for a total of 13-26 cell types with technical triplicates per day. This technology enabled us to dissect the complex spatial relationships among immune cells, tumor cells, and the surrounding vasculature, providing a detailed view of the TME and its proteomic heterogeneity.

To minimize potential proteomic alterations due to the cyclic multiplexed imaging process, which involves multiple rounds of staining and imaging, we used photobleaching instead of chemical stripping. The MACSima facilitated direct and automated antibody pipetting onto membrane slides within a hub, reducing slide handling between imaging cycles. To evaluate the impact of MACSima multiplex staining on proteome integrity, we isolated tissue areas of 40,000 µm², equivalent to approximately 60 CRC cells, from both immunofluorescence-unstained control and MACSima-stained CRC tissues (**Figure 1D**). Comparative analysis of the precursor and identified protein profiles revealed only minimal protein loss during the staining process (no loss of proteins identifications after 14-plex), confirming the compatibility of MACSima imaging with downstream laser microdissection and the efficiency of protein recovery for MS-based proteomics (**Figure 1D**). Similarly, we observed minimal protein loss during the staining process for tonsil cancer tissues (less than 2% loss of protein identifications after 22-plex staining) (**Figure S1B**), demonstrating the broad applicability of this approach across different tumor types.

We conclude that the integration of high-plex MACSima imaging, automated single-cell microdissection, and MS-based proteomics enables in-depth, spatially resolved molecular characterization of the complex TME. This multimodal approach preserves the spatial context of the tissue while achieving deep proteome coverage, offering new opportunities to decipher the intricate cellular interactions and signaling networks that shape tumor progression and response to therapy.

### Macrophages establish an immunosuppressive barrier hindering T cell infiltration in colorectal cancer

To investigate the TME with respect to immunologically hot or cold properties, we conducted 14-plex staining on a CRC tumor (**Experimental Methods**). We identified distinct cell populations within the CRC tissue, including tumor cells, macrophages, cytotoxic T cells (CTLs), helper T cells (TH), B cells, granulocytes, as well as blood vessels and lymphatic vessels. The presence and spatial distribution of these cell types are crucial for understanding the immunological landscape of the tumor, which in turn informs the choice of immunotherapy (**Figure 1B**; **Figure 2A; Figure S1C**).

**Figure 2.**
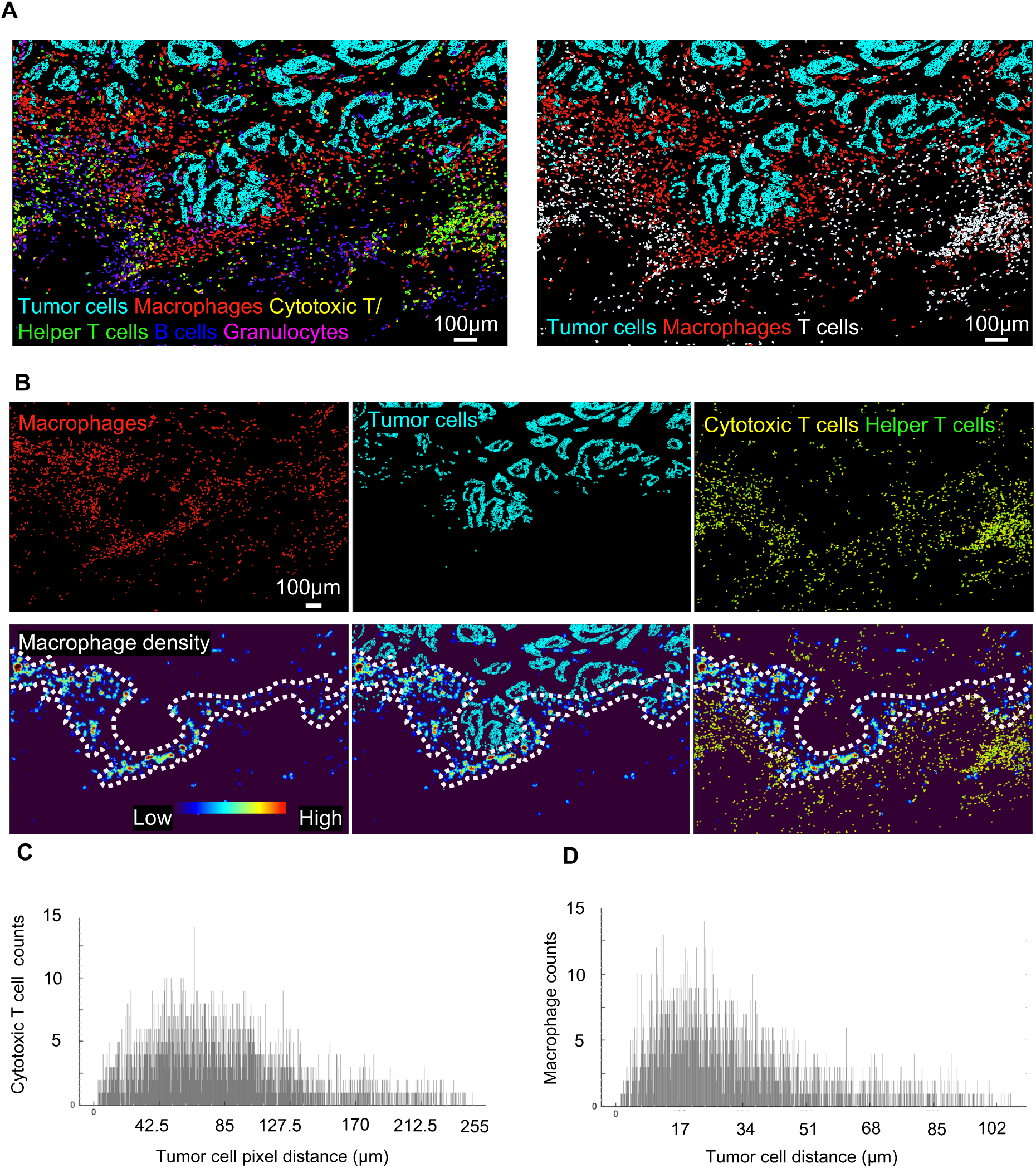
Macrophages establish an immunosuppressive barrier hindering T cell infiltration in colorectal cancer. *(A) Phenotype map showing the spatial distribution of the indicated cell types (tumor cells, cytotoxic T cells, helper T cells, B cells, granulocytes and macrophages) in the tumor microenvironment (TME). (B) Representative image of a macrophage-abundant region with overlaid masks highlighting tumor cells and T cells (cytotoxic and helper T cells). The barrier-like positioning of macrophages between the tumor epithelium and lamina propria suggests their role in shaping the immune landscape. (C-D) Histogram of the counts of cytotoxic T cells (C) or macrophages (D) as a function of their proximity from tumor cells in colorectal cancer, documenting the spatial relationship between these cell types*.

The image analysis yielded two main findings: firstly, we observed a remarkable accumulation of immune cells within the lamina propria, positioned beneath the tumor epithelium (**Figure 2A**). Secondly, we noted a barrier-like positioning of macrophages between the tumor epithelium and lamina propria, characterized by low expression of HLA-DR (**Figure 2B, Figure S1D**), suggesting their reduced antigen-presenting capacity and involvement in immunosuppression ^26^. Additionally, we observed a higher abundance of immune cells, including CTLs and helper T cells, infiltrating the lamina propria compared to the tumor epithelial region (**Figure 2B**).

Spatial distribution analysis revealed proximity of macrophages (peak count at distance of 68 μm) to the tumor epithelium compared to CTLs (peak count at distance of 20 μm). This indicates the formation of an immunosuppressive barrier by macrophages that hinder the infiltration and activity of T cells within the tumor epithelium ^27^, thereby contributing to an immunologically cold TME (**Figure 2C-D**). The presence of at least a few T-cells in the tumor epithelial region further classifies it as ‘T-cell excluded’ rather than ‘immune desert’ ^28^. The distinct infiltration pattern of T cells in the lamina propria and tumor epithelium, separated by a barrier formed by macrophages, suggested the need for future proteome analysis of T cells in both regions, elucidating functional differences and interactions between immune cell subsets.

### T cell subsets have distinctive spatial proteome signatures in colorectal TME

To functionally characterize T-cells in a spatial context, we conducted single-cell microdissection of CTLs and TH from both tumor epithelial and lamina propria regions, followed by deep proteome profiling. Our analysis quantified more than 4000 protein groups in both CTLs and TH from the equivalent of only 200 of these very small cells. This revealed substantial differences in protein expression profiles for the same cell types between the two regions (**Figure 3A-B**). Interestingly, unbiased proteomics revealed the expression levels of markers of cytotoxic T cells (including CD3, CD8, perforins and granzymes) and helper T cells (including CD3, FOXP3 and IRF4). Overall, more than 1000 proteins were significantly differentially expressed proteins in CTLs between the lamina propria and tumor epithelial regions (**Figure 3C**), and about 200 in TH between the same regions (**Figure 3D**).

**Figure 3.**
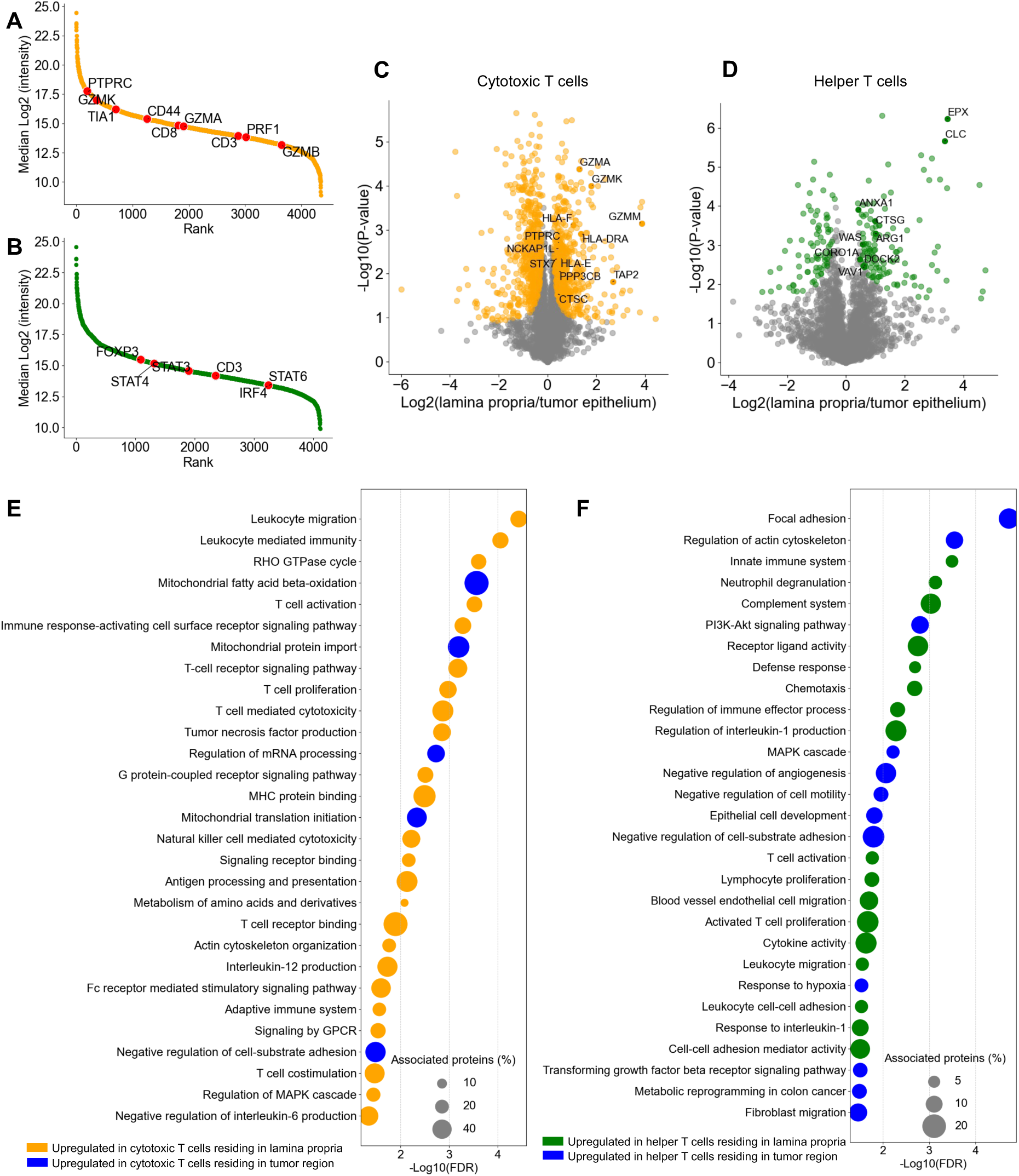
T cell subsets have distinctive spatial proteome signatures in colorectal tumor microenvironment. *(A-B) Ranking of protein quantified in cytotoxic T cells (CTLs) (A) and helper T cells (TH) (B), ranked by median transformed intensity values detected by mass spectrometry, with cell-type specific markers highlighted. (C-D) Volcano plots illustrating proteomic comparisons of CTLs (C) and TH cells (D) between the lamina propria and tumor epithelial regions, with T-cell activity-related markers highlighted. Significantly enriched proteins are denoted as red dots (two-sided t-test, FDR < 0.05, S0 = 0.1). (E-F) Gene Ontology (GO) term analysis of significantly altered proteins in CTLs (E) and TH (F) residing in the lamina propria and tumor epithelial regions, revealing functional differences between T cell subsets in distinct tumor compartments*.

Comparative analysis of the quantitative proteomes of CTLs revealed distinct functional characteristics between those residing in the lamina propria and tumor epithelium. Pathways associated with T cell activation, cytokine production, immune receptor signaling, and antigen processing and presentation were down-regulated in CTLs residing in the tumor epithelial region, indicating reduced immune response against cancer cells (**Figure 3E**). In contrast, in the laminar propria, pathways related to cytotoxicity, such as natural killer cell-mediated cytotoxicity, T-cell-mediated cytotoxicity, MHC protein binding, tumor necrosis factor production and leukocyte-mediated immunity were elevated, suggesting increased effectiveness in directly immune response. These CTLs also had comparatively upregulated MAPK cascade, GPCR signaling, interleukine-12 production and T cell receptor signaling pathway, implying enhanced cellular communication and activation of intracellular signaling cascades crucial for immune response regulation. Their effective functioning appears to be supported by elevated metabolic pathways. Moreover, upregulation of cell migration and adhesion suggest active movement within tissues to locate and engage with cancer cells. Conversely, CTLs in the tumor epithelium exhibited upregulated mitochondrial metabolism including mitochondrial fatty acid beta-oxidation to adapt to the TME. Additionally, MS-based proteomics pinpoints the identity and fold-change of the involved molecular players. For instance, molecules such as CTSC, GZMM, GZMA, GZMK, HLA-DRA, HLA-E, HLA-F, NCKAP1L, PPP3CB, PTPRC, STX7, and TAP2, which play crucial roles in T cell cytotoxicity, exhibit significant upregulation in cytotoxic T cells located in the lamina propria, with fold changes ranging between 2 and 16 (**Figure 3C**).

Similarly, the proteomics analysis of TH in the tumor epithelial region revealed distinct functional roles compared to those in the lamina propria. In the tumor epithelium pathways associated with adaptation to hypoxic TME, metabolic reprogramming, negative regulation of angiogenesis, and cell adhesion were upregulated, suggesting their involvement in modulating immune responses, tumor progression, and interactions with the TME (**Figure 3F**). Furthermore, growth factor signaling such as the transforming growth factor beta receptor (TGFRb) signaling pathway, and intracellular signaling cascades including PI3K-Akt signaling and MAPK cascade, indicated alterations in responsiveness to external stimuli. These changes may enable TH in the tumor epithelium to actively participate in tumor cell proliferation, survival, and immune evasion mechanisms. Upregulated pathways in lamina propria-residing TH included T cell activation, interleukin-1 response and monocyte chemotaxis. This suggests increased antigen processing and presentation capability and increased immune surveillance. The activation of pathways linked to the innate immune system, such as modulation of the complement system and neutrophil degranulation, alongside elevated cell-cell adhesion, underscores the significant role of lamina propria-residing TH in modulating immune responses, immune cell localization, and communication. Molecules such as ANXA1, ARG1, CLC, CORO1A, CTSG, DOCK2, EPX, VAV1, WAS, which play crucial roles in T cell activation, exhibit significant upregulation in TH located in the lamina propria, with fold changes ranging between 1.5 and 16 (**Figure 3D**).

These findings highlight the distinct functional roles of CTLs and TH residing in different microenvironments within CRC directly at the level of the proteins involved. Our data provides detailed insights into the interactions with the TME with potential implications for tumor progression and immune response modulation.

### Immune cells abundantly infiltrate and cytotoxic T cells closely position in tonsil cancer

To evaluate the versatility of our spatial proteomics technology, we additionally extended it to tonsil cancer FFPE sections, incorporating expanded marker staining (22-plex) to enhance the detection of a more diverse range of cell types (**Figure 4A; Figure S1A**). Given our extensive range of markers, we were able to map intercellular spatial relationships between cell types enabling investigation of the interplay between cells in the TME. We assessed the expression of Cytokeratin (CK), CD3, CD4, CD8, CD20, CD38, CD57, CD11b, CD68, CD31, Podoplanin, HLA-DR and Ki67, based on the staining and performed Uniform Manifold Approximation and Projection of 57,276 cells. We identified four primary cell clusters: tumor cells, lymphocytes, myeloid cells, and endothelial cells. In-depth examination of lymphocytes further revealed distinct populations, comprising cytotoxic T cells, helper T cells, B cells, plasma cells, and natural killer (NK) cells (**Figure 4B-C**). Notably, immune cells constituted about 43% of the total identified population in the tumor (**Figure 4D**), highlighting substantial immune cell infiltration.

**Figure 4.**
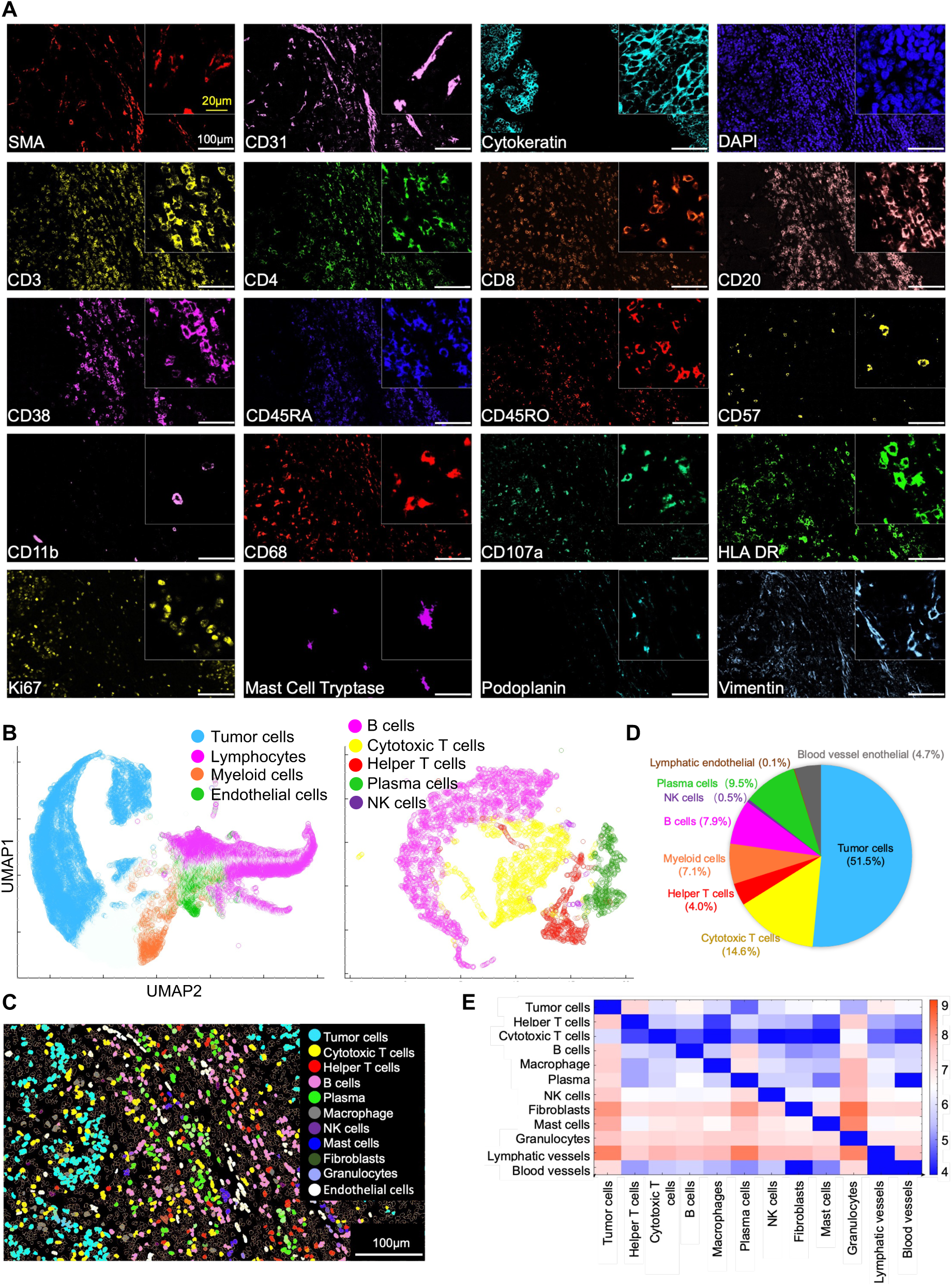
Immune cells abundantly infiltrate and cytotoxic T cells closely position in tonsil cancer. *(A) Representative images: multiplex staining on tonsil cancer tissue sections. Multiplexed pseudo-colored images display marker staining within the same tissue region, providing a comprehensive view of the cellular composition. Scale bar (in white), 100 µm. (B) Cell types: UMAP revealing four distinct clusters. The lymphocyte cluster further divides into five subclusters, based on the expression of CK, CD3, CD4, CD8, CD20, CD38, CD57, CD11b, CD68, CD31, Podoplanin, HLA-DR and Ki67. (C) Identified, colored masks represent cellular classification in a representative region, visualizing the spatial distribution of different cell types. (D) Tumor cells constitute about half of classified cell types, with immune cells as the second largest category (43%), whereas the remainder are endothelial cells. (E) Distance heatmap depicting the proximity and interactions among classified cell types. Pixel distances among individual cells were arcsinh normalized and averaged, with lower values indicating higher proximity (**Experimental Methods**)*.

To gain insights into the spatial organization of the TME, we evaluated the spatial distances among diverse cellular populations. To do so, we determined all pairwise pixel distances between different cell types, averaged and normalized them and constructed a heat map (**Experimental Methods**, **Figure 4E**). This analysis revealed that CTLs were positioned closely to tumor cells, with an average normalized distance value of 6 (Arcsinh normalized and averaged pixel distances among individual cells, ranging from 4 to 9, indicating closer to farther distance, respectively). In comparison, the distance values within the same cell type were the closest at 4, while distances between tumor cells and other cell types such as helper TH, B cells, macrophages, and natural killer (NK) cells ranged from 7 to 9, indicating greater distance. Furthermore, CTLs were found to be relatively close to other stromal cells, including helper T cells, B cells, macrophages, plasma cells, NK cells, endothelial cells, and fibroblasts, with average normalized distances ranging from 5.5 to 6.5. This proximity suggests significant infiltration of CTLs into both the tumor parenchyma and stroma, potentially facilitating interactions or crosstalk between CTLs and tumor cells, indicating that this TME is immunologically hot ^19^.

The close positioning of CTLs to tumor cells in tonsil cancer, as revealed by our spatial analysis, reveals where direct tumor-immune interactions and anti-tumor immune responses take place. The abundant immune cell infiltration and close proximity of CTLs to tumor cells in tonsil cancer suggest a more permissive TME for anti-tumor immunity, which may contribute to the immunologically hot nature of this cancer type. This physical proximity is in stark contrast to the immunosuppressive macrophage barrier observed in colorectal cancer, which hinders T cell infiltration into the tumor epithelium. These insights highlight the importance of considering the spatial context and cellular interactions when assessing the immunological status of a tumor and its potential responsiveness to immunotherapies.

### CTLs infiltrate to tumor parenchyma with a suppressive impact on tumor proliferation

We found that CTLs not only abound in the stroma but also significantly infiltrate the tumor parenchyma, whereas TH primarily reside in the stroma (**Figure 5A**). This heightened proximity of CTLs to tumor cells as compared to TH indicates their potential role in tumor surveillance and anti-tumor response.

**Figure 5.**
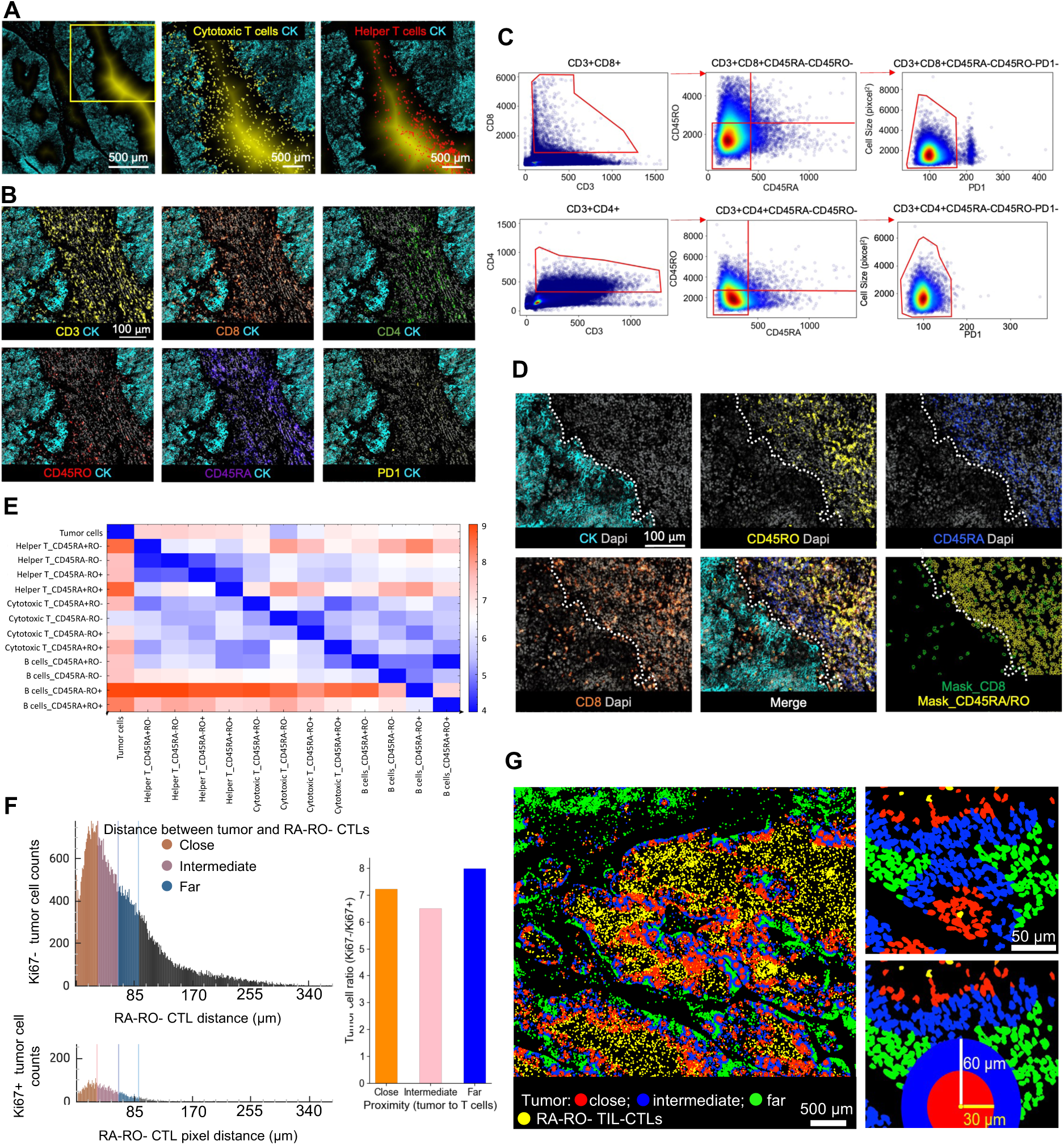
Cytotoxic T cells (CTLs) infiltrate to tumor parenchyma with a suppressive impact on tumor proliferation. *(A) Density gradient in the background indicates distance to tumor islands, with brighter yellow representing greater distance from tumor cells. Left panel only has the gradient whereas the middle, zoomed panel additionally depicts yellow masks of the cytotoxic T cells and the right panel red dots of helper T cells, respectively. (B) Representative images showcasing markers for T cell phenotyping, enabling the identification and characterization of different T cell subsets. (C) T cell gating based on the expression of CD3, CD8, CD45RA, CD45RO, and PD1 to analyze the corresponding T-cell subpopulations. (D) Representative images showing tumor-infiltrating CD3 and CD8 positive CTLs with absent CD45RA and CD45RO expression. (E) Distance heatmap depicting proximity and interactions between tumor cells and T and B lymphocyte subtypes, considering CD45RA and CD45RO expression. Distances among individual cells were arcsinh normalized and averaged, with lower values indicating higher proximity as in* Fig. 4E*. Note that CD45RA-CD45RO- (RA-RO-) CTLs are the closest to tumor cells. (F) Histogram of Ki67-/Ki67+ tumor cell number as a function of pixel distance from RA-RO-CTLs. Ki67-/Ki67+ tumor cell ratios were quantified within the same imaged region. (G) Representative image displaying masks of tumor cells at varying proximities to RA-RO-tumor-infiltrating CTLs, enabling subsequent isolation and proteomic analysis of tumor cells based on their spatial relationship to TIL-CTLs*.

CD45RA and CD45RO, transmembrane glycoproteins found on leukocytes, are markers for resting T cells and memory T cells activated by antigens, respectively ^29^. Our analysis identified two primary subpopulations: CD45RA-CD45RO- (abbreviated as RA-RO-) as the most prevalent, followed by CD45RA-CD45RO+ (abbreviated as RA-RO+) (**Figure 5B-C**). Notably, both CTLs and TH within the RA-RO-subgroup exhibited a lack of PD1 expression (**Figure 5C**).

In the tumor-infiltrating cytotoxic T lymphocytes (TIL-CTLs) we unexpectedly found a new phenotype, namely TIL-CTLs that had lost expression of CD45RA and CD45RO (**Figure 5B-D**). The observed diminished CD45RA or CD45RO expression in TIL-CTLs suggests a specialized functional adaptation within the TME, warranting further investigation (**Figure 5D**). Among all CD45RA and CD45RO-defined lymphocyte subpopulations, RA-RO-CTLs were the closest type in proximity to tumor cells, further supporting the notion that TIL-CTLs are RA-RO- (**Figure 5E**).

To assess the functional impact of TIL-CTLs on tumor cells, we investigated the relationship between TIL-CTL proximity and tumor cell proliferation. Remarkably, we observed a suppressive effect of TIL-CTLs on tumor cell proliferation, as evidenced by a higher ratio of Ki67-/Ki67+ tumor cells (indicating reduced proliferation) in tumor cells located within close proximity (<30 µm) to TIL-CTLs compared to those at intermediate distances (>30 µm and <60 µm) **(Figure 5F).** This finding aligns with the observed upregulation of the degranulation marker CD107 on CTLs, a critical factor for enhanced cytotoxic activity 30 **(Figure S2A)**. Unexpectedly, tumor cells positioned even further away (>60 µm and <90 µm) from TIL-CTLs had a more pronounced increase in the Ki67-/Ki67+ ratio (**Figure 5F**). This suggested heightened immune surveillance in regions where tumor cells are more distant from TIL-CTLs and closer to the stroma, with potential implications for regulating tumor growth.

To further investigate this unexpected phenomenon and gain insights into the underlying molecular mechanisms, we performed spatial and cell type-specific unbiased proteomics. To this end, we isolated tumor cells according to their proximity to TIL-CTLs, categorizing them as close, intermediate, and far, for subsequent proteome analysis. The premise of our strategy was that the proteomic diversity of tumor cells based on their distance from TIL-CTLs, might offer insights into the molecular landscape and potential functional consequences of these distinct tumor cell populations (**Figure 5G**).

### Proteomic heterogeneity in tumor cells correlates with proximity to TIL-CTLs

Tumor cells located at various proximities — close (<30 µm), intermediate (>30 µm and <60 µm), and distant (>60 µm and <90 µm) — to TIL-CTLs exhibited different proteomic signatures (**Figure 6A-B**). Principal Component Analysis (PCA) revealed distinct grouping of these tumor cells according to their proximity to TIL-CTLs, with those in the distant group clustering notably apart from the close and intermediate groups (**Figure 6B**).

**Figure 6.**
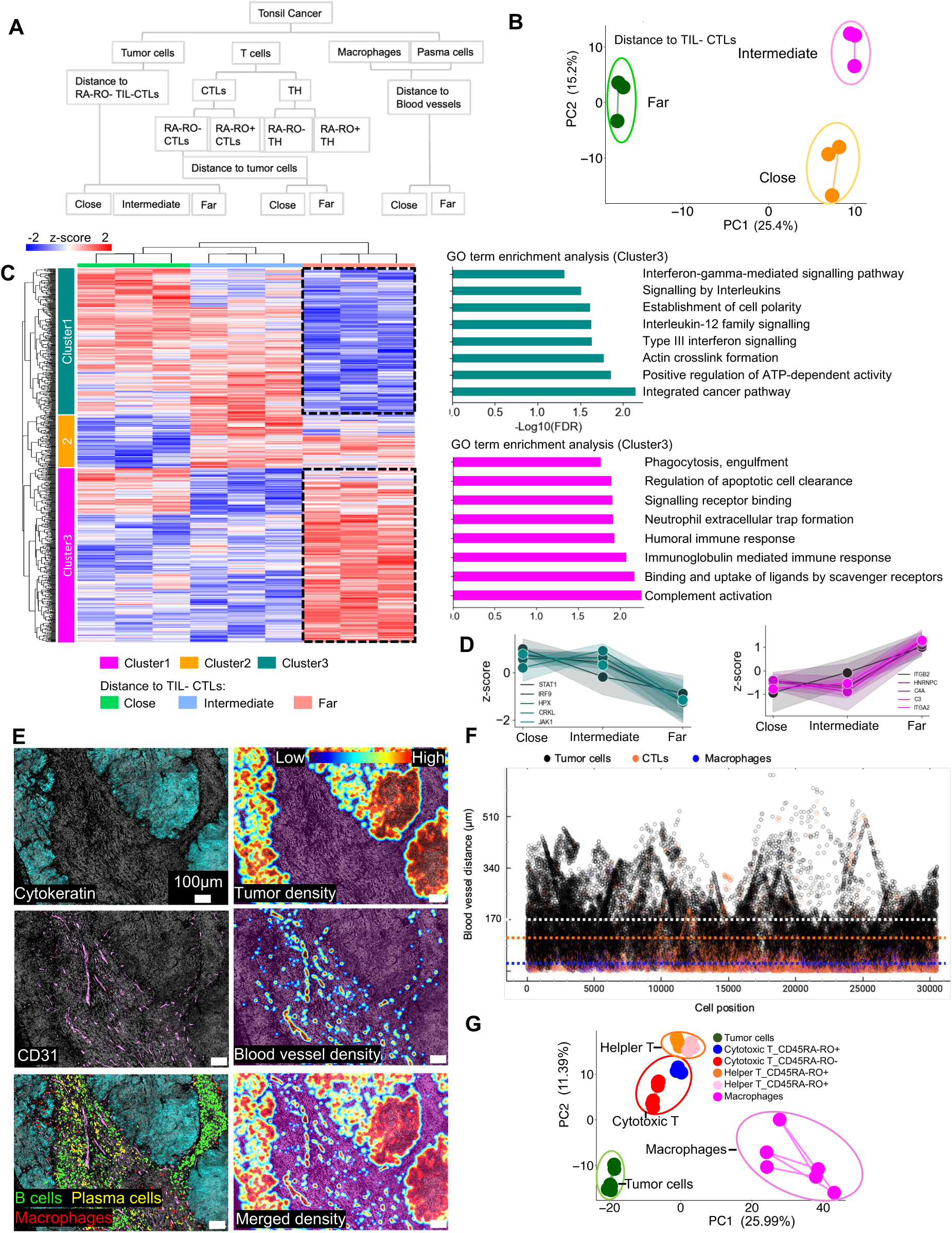
Proteomic heterogeneity in tumor cells correlates with proximity to tumor-infiltrating lymphocyte cytotoxic T cells (TIL-CTLs). *(A) Diagram illustrating the spatial localization of cells isolated from tonsil cancer tissues for proteome profiling, enabling the investigation of proteomic heterogeneity based on cellular proximity. (B) Principal component analysis (PCA) of tumor cell proteomes reveals distinct clustering based on proximity to TIL-CTLs. (C) Heatmap highlighting significantly differentially expressed proteins among tumor cells with different proximity to TIL-CTLs (686 ANOVA significant proteins, FDR < 0.05). Gene Ontology (GO) term analysis of indicated clusters is shown, providing insights into the functional implications of the proteomic heterogeneity. (D) Profile plots of key molecules in interferon, interleukin-12 signaling pathways, and phagocytic processes. (E) Representative images illustrating tumor islands, blood vessels, and the distribution of B cells, plasma cells, and macrophages. The colored areas represent tumor or blood vessel densities, with a color gradient from red (highest density) to blue (lower density at the edges). This visualization aids in understanding the spatial relationships between different cellular components. (F) Dot plot displaying the distance of tumor cells, TIL-CTLs, and macrophages to blood vessels. The white dotted line represents the median distance between tumor cells and blood vessels, the blue dotted line shows the median distance between TIL-CTLs and blood vessels, and the orange dotted line marks the median distance between macrophages and blood vessels. (G) PCA of proteomes of tumor cells, lymphocytes, and macrophages, demonstrating distinct clustering based on cell type and highlighting the proteomic differences between these cellular populations*.

The upregulated interferon and interleukin-12 signaling in tumor cells close to TIL-CTLs indicate that tumor cells response to cytotoxic activity of CTLs and Th1 (**Figures 6C**). The key players involved in this include JAK1, STAT1, IRF9, HPX, and CRKL (**Figures 6D**). Conversely, tumor cells distant from TIL-CTLs and in proximity to stroma exhibited a higher Ki67-/Ki67+ tumor cell ratio compared to those in closer proximity (**Figure 5F**), which was corroborated by increased pathway activity in complement activation, immunoglobulin-mediated immune response, and phagocytosis engulfment in these distant tumor cells (**Figures 5C**). This suggests active immune surveillance, potentially from macrophages, B cells, and plasma cells in the stroma. Proteins involved in this response include C3, C4A, ITGA2 and ITGA2, are known to increase the interaction between macrophages and tumor cells, facilitating phagocytosis^30^ (**Figures 6D**). The high density of macrophages, B cells, and plasma cells in the tumor stroma may indicate physical interactions between them and tumor cells at the stromal edge (**Figures 6E**). Blood vessels were predominantly present within the tumor stroma, making the distance of various cell types to these vessels a reliable indicator of their closeness to the tumor stroma itself (**Figures 6E**).

To assess the spatial distribution across the entire imaged area and to present this information through dimensionality reduction techniques, we charted the x-axis positions of tumor cells, macrophages, and CTLs relative to their pixel distance from blood vessels. A higher pixel distance value indicates a greater distance of tumor cells, macrophages, and CTLs from the blood vessels located in the stroma. The median distance from blood vessels to tumor cells was 170 μm, whereas this was 119 μm to CTLs and 17 μm to macrophages. Thus, on the basis of physical distance, tumor cells far from blood vessels could be attacked by CTLs, while those closer to vessels also by macrophages (**Figure 6F**).

To investigate this on a molecular basis, we isolated macrophages from areas close and far from blood vessels for proteome analysis (**Figure 6F**). We set a cutoff at 85 µm to distinguish between close and far distances to blood vessels, based on it being half of the median distance observed between tumor cells and blood vessels (**Figure 6F**). PCA analysis revealed distinct grouping of tumor cells, T cells, and macrophages (**Figure 6G**). The notable upregulation of membrane dynamics regulators such as ACAP1 in macrophages near blood vessels suggests increased cellular mobility, aiding their migration from blood vessels to tumor sites ^31^ (**Figure S2B**). These findings, consistent with image analysis, underscore the functional interplay and cellular dynamics between the immune system and tumor cells within the tumor parenchyma and its immune-enriched stroma.

### TIL-CTLs exhibit strategic adaptations to hypoxic tumor microenvironment

Following our analysis of how the spatial proximity to TIL-CTLs impacts the functional state of tumor cells as judged by their proteomes, including respond to the cytotoxic activity of CTLs, we next explored how TIL-CTLs themselves undergo proteomic alterations as a function of the TME. As described for CRC above, we first obtained proteomic profiles for the most abundant TIL-CTL subpopulations: CD45RA-CD45RO- (RA-RO-) and CD45RA-CD45RO+ (RA-RO+) CTLs (**Figure S2C**). Between these subpopulations, there were 177 significantly regulated proteins. RA-RO+ CTLs exhibited higher cytotoxicity as judged by pathway analysis (**Figure 7A-B**). Additionally, RA-RO-CTLs adapted to the TME strategically, as indicated by upregulated response to hypoxia (including NRF2). Compared to their RA-RO+ counterparts residing in stroma, this would facilitate their penetration into deeper tumor regions, more hypoxic tumor region ^32^ (**Figure 7A-B**). Consistent with this, molecules such as ANXA1, GSTP1, PCNA, TRAP1 and TXN, which play crucial roles in hypoxia adaptation ^33,34^, were significantly upregulated in RA-RO-CTLs, with fold changes ranging between 1.5 and 4 (**Figure 7A**).

**Figure 7.**
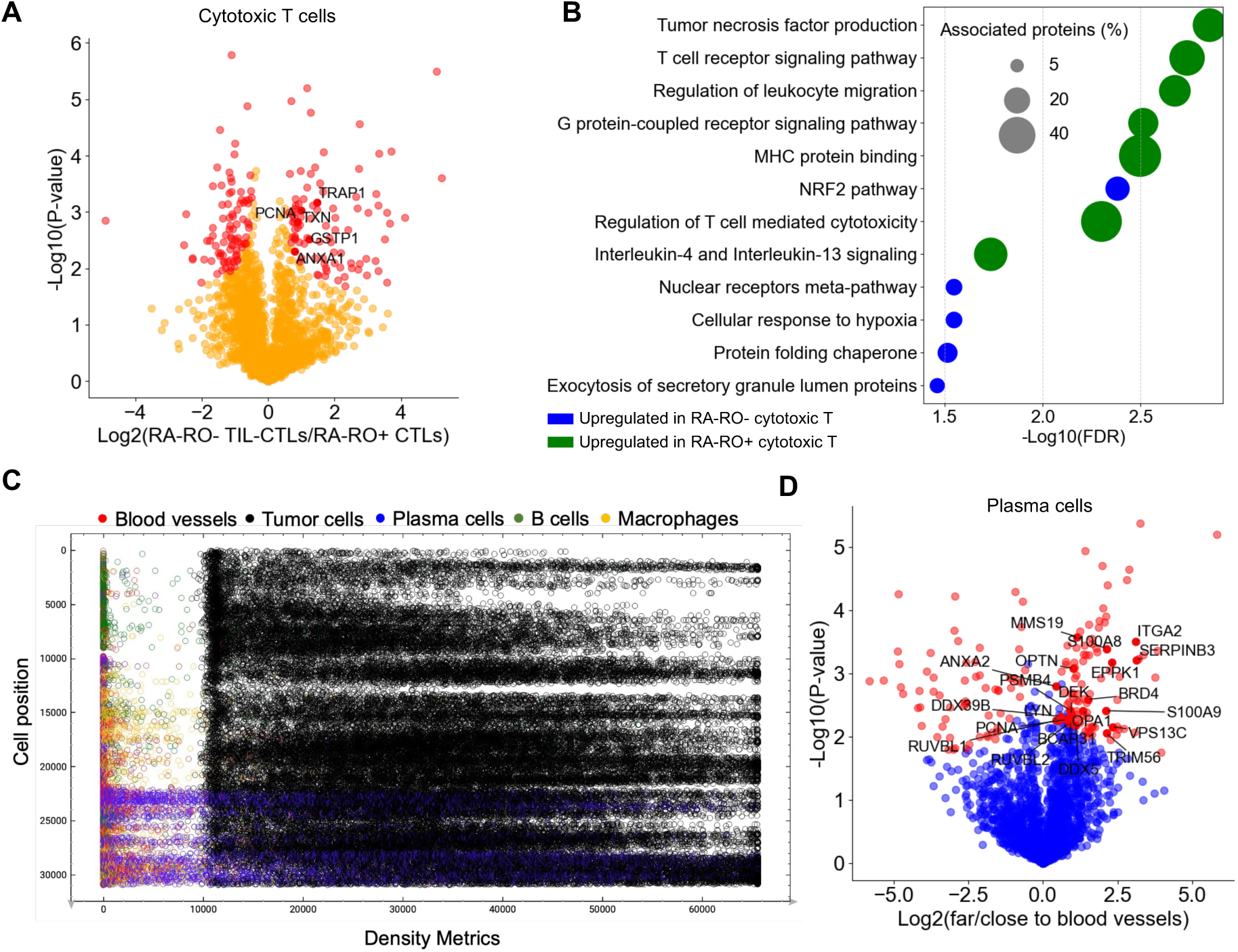
Tumor-infiltrating lymphocyte cytotoxic T cells (TIL-CTLs) exhibit strategic adaptations to hypoxic tumor microenvironment. *(A) Volcano plot depicting pairwise proteomic comparisons between CD45RA-CD45RO- (RA-RO-) CTLs and RA-RO+ CTLs. Significantly enriched proteins are shown as red dots (two-sided t-test, FDR < 0.05, S0 = 0.1). (B) Gene Ontology (GO) term analysis of significantly altered proteins in RA-RO-TIL-CTLs compared to their RA-RO+ counterparts, highlighting the upregulation of pathways related to hypoxia response and NRF2 signaling. These adaptations facilitate TIL-CTL infiltration into the hypoxic tumor microenvironment. (C) Density metrics displaying cell type densities relative to nuclear X position, with higher values signifying greater density. (D) Volcano plot depicting pairwise proteomic comparisons between plasma cells in regions far from and close to blood vessels. Significantly enriched proteins are highlighted as red dots (two-sided t-test, FDR < 0.05, S0 = 0.1)*.

In addition to the cytotoxic T cells and macrophages investigated above, other cell types are involved in immune surveillance of tumor cells including plasma cells, which are plentiful in the stroma (**Figures 6E**). To visualize the spatial distribution of these cell types, we plotted cellular density metrics along the x-axis position, where a higher density metrics value indicates a greater density at that position. This visualization effectively distinguished blood vessels (with density metrics from 0 to 10,000) from tumor cells (with density metrics from 10,000 to 65,000), accurately representing their spatial density (**Figure S2D**). We extended our analysis to encompass tumor cells, blood vessels, B cells and plasma cells, which revealed widespread distribution from vessel-rich areas to tumor sites (**Figure 7C**). Therefore, to explore proteomic differences in immune cells between vessel-rich areas and tumor sites, we isolated immune populations based on their proximity to blood vessels—those close to (<85 µm) and those far from (>85 µm) blood vessels, corresponding to being near and distant from tumor sites, respectively. We established a cutoff at 85 μm because this was half of the median distance between tumor cells and blood vessels (**Figure 6F**).

The CTLs proximal to tumor cells exhibited downregulation of LAMA1/4, TNC, and ITGA6 compared to those distal from tumor cells, suggesting enhanced T cell mobility and improved penetration into the tumor parenchyma ^35,36^ (**Figure S2E**). Plasma cells located distant from vessels and closer to the tumor upregulated the stress responses, interleukin-3/5, and GM-CSF, along with increased chemotaxis response. This suggests active migration towards the tumor site and adaptation to the stress of the TME for effective immune surveillance. Conversely, plasma cells positioned near vessels and away from the tumor core upregulated pathways associated with RHO GTPase effectors, calcium ion response, and actin cytoskeleton organization, perhaps indicating strategic motility adaptation within the TME ^37^ (**Figure 7D**; **Figure S2F**). Together, these findings highlight the dynamic nature of immune responses and the adaptive capabilities of CTLs and plasma cells in the TME, emphasizing the importance of spatial context in understanding the functional states of immune cells within the complex tumor ecosystem.

## Discussion

In this study, we introduce an innovative approach that integrates highly multiplexed imaging with deep visual proteomics (DVP), enabling the spatial profiling of distinct cell populations within the tumor microenvironment (TME) at an unprecedented level of detail. To achieve this we had to solve the challenges with multiplex staining on membrane slides related to the delicate nature of membrane slides and their incompatibility with heating or chemical stripping steps, leading to protein loss. We employed automated staining, imaging and photobleaching cycles, enabled us to maintain tissue and protein integrity. This novel integration of technologies allows us to uncover intricate tumor-immune interactions and functional adaptations, providing new insights into cancer biology and revealing potential targets for immunotherapy.

Our approach seamlessly combines spatial information from multiplexed imaging with the unbiased, high-depth protein discovery capabilities of DVP. This integration surpasses the limitations of existing methods by enabling the visualization and quantification of thousands of proteins within the TME, offering a comprehensive view of the complex interplay between tumor and immune cells at a functional level.

We applied this approach to colorectal cancer (CRC) and tonsil cancer samples, revealing significant differences in the proteome signatures of cytotoxic T lymphocytes (CTLs) and helper T cells (TH) between the lamina propria and tumor epithelium in CRC. These findings highlight the impact of the microenvironment on immune cell functional states. Notably, we identified an immunosuppressive macrophage barrier in CRC that impedes T cell infiltration into the tumor epithelium. It may be mediated by cell-cell adhesion, secretion of extracellular matrix components, or establishment of chemotactic gradients ^27,38^. Targeting the macrophages in the barrier could be a promising therapeutic strategy to enhance T cell-mediated anti-tumor immunity, potentially by disrupting the physical or chemical signals that restrict T cell infiltration ^38^. suggesting potential therapeutic strategies to enhance T cell-mediated anti-tumor immunity.

In tonsil cancer, our 22-plex imaging revealed substantial immune cell infiltration, with CTLs closely positioned to tumor cells and infiltrating both the parenchyma and stroma. Tumor-infiltrating CTLs exhibited a loss of CD45RA and CD45RO expression, indicating specialized functional adaptations within the TME. Our proteomic analysis showed that these CD45RA-CD45RO- (RA-RO-) TIL-CTLs upregulated pathways related to hypoxia response and NRF2 signaling, facilitating their penetration into hypoxic tumor regions.

Furthermore, our spatial proteomics approach uncovered significant tumor cell proteomic heterogeneity influenced by proximity to TIL-CTLs. Tumor cells distant from TIL-CTLs showed increased activity in complement activation and phagocytosis, suggesting active immune surveillance from macrophages, B cells, and plasma cells in the stroma. These insights highlight the importance of considering the spatial context and cellular interactions when assessing the immunological status of a tumor and its responsiveness to immunotherapies.

Our integrated approach offers several advantages over existing methods: it provides a detailed view of the TME by combining spatial information with deep proteomic data, enables the discovery of novel proteins and pathways that may be overlooked by targeted approaches, and allows for a better understanding of the complex interactions between tumor and immune cells. As we refine and apply this approach to a broader range of cancer types and immunotherapy strategies, we anticipate that it will play a pivotal role in advancing our understanding of tumor biology and guiding the development of personalized cancer therapies. Furthermore, our approach could help identify the optimal spatial positioning and functional state of engineered T cells, such as chimeric antigen receptor T (CAR-T) cells and T cell receptor-engineered T (TCR-T) cells, within the TME, providing valuable insights into factors influencing their efficacy and guiding strategies to improve their performance ^39^.

While our study provides novel insights into the TME, further validation of the identified cellular adaptations and interactions using functional assays would provide a more comprehensive understanding of their biological significance. By enabling the precise profiling of an individual patient’s TME, this approach holds great promise for transforming patient care, allowing for tailored treatment strategies and monitoring of responses.

## Materials and Methods

### Tissue specimen preparation

FFPE tissue specimens of human tonsil cancer were procured from ProteoGenex (ProteoGenex, USA). The patient, a 48-year-old male, was diagnosed with stage IV primary tonsil squamous cell carcinoma. Colorectal cancer FFPE tissue was obtained from Zealand University Hospital in Denmark and fully anonymized. All specimens were acquired with informed consent, following standard protocols, and received appropriate approval by Health Research Ethics Approval (VEK) and ProteoGenex. Membrane PEN slides 1.0 (415190-9041-000, Zeiss) were exposed to UV light for 1 hour and coated with Vectabond reagent (SP-1800-7, Vector Laboratories) following the manufacturer’s protocol. FFPE tissue sections, cut at a thickness of 3 µm, were mounted on treated PEN slides, left to dry at 37°C overnight, and heated at 60°C for 20 minutes to enhance tissue adhesion ^15^. Subsequently, paraffin removal and hydration were performed by sequential passage through xylol and decreasing concentrations of ethanol (EtOH) as follows: xylenes (3 × 5 min), 100% EtOH (2 × 2 min), 95% EtOH (2 min), 80% EtOH (2 min), 70% EtOH (2 min), and distilled and deionized water (ddH_2_O, 2 min). Antigen retrieval utilized EDTA buffer (pH 8.5, E1161, Sigma) and 10% glycerol (G7757, Sigma Aldrich), heated for 45 minutes at 88°C in a water bath. Thereafter, the two-well MACSwell Imaging Frame was promptly mounted onto the slide, after which 550 µL MACSima Running Buffer was added.

### MACSima multiplex-staining and imaging

The MACSima Imaging System by Miltenyi Biotec is a fully automated instrument seamlessly integrating liquid handling with widefield microscopy for cyclic immunofluorescence imaging^21^. Prior to initiating the MACSima instrument, a DAPI pre-staining procedure was executed: The MACSima Running Buffer was extracted from the target sample, which was then stained for 10 minutes with a 1:10 dilution of a DAPI staining solution. Following staining, three washing steps were conducted using MACSima Running Buffer, and ultimately, 550 µL of MACSima Running Buffer was added. The antibodies employed in the current study were procured from Miltenyi Biotec and are enumerated below: CD3-APC (130-120-269), FOXP3-PE (130-127-808), Mast Cell Tryptase-APC (130-125-278), PD1-PE (130-117-384), CD107a-APC (130-126-203), CD57-PE (130-111-810), CD38-PE (130-126-438), HLA-DR-FITC (130-123-076), CD4-PE (130-127-906), CD45RA-APC (130-112-097), CD20-FITC (130-118-292), CD8-PE (130-117-201), CD45RO-APC (130-113-556), Ki67-FITC (130-117-691), Cytokeratin-FITC (130-112-743), CD11b-PE (130-128-773), Podoplanin-APC (130-126-165), CD68-PE (130-128-345), Vimentin-FITC (130-116-508), SMA-FITC (130-123-363), CD31-PE (130-128-769). Unspecific binding sites were blocked using MACSima FCR Block. Hybrid autofocusing was achieved through a combination of hardware autofocus and image-based autofocusing optimized through the DAPI image. Imaging encompassed an exposure series of 6, 36, 216 ms in the APC channel, 5, 20, 80 ms in the PE channel, and 8, 32, 128 ms in the FITC channel. Photobleaching was employed to eliminate the signal. Image acquisition utilized an epifluorescence setup with a long-working-distance 20 × objective (NA 0.45, objective slides 1.0 mm).

### Image analysis and cell extraction

The MACS iQ View Software facilitated image preprocessing workflow ^21^. This involved correcting image distortion, stitching, registering images for spatial alignment, and subtracting residual intensities resulting from remaining autofluorescence or incomplete erasure of the previous cycle staining. Image segmentation was achieved using nuclei and cell membrane markers. A gating-based strategy facilitated cellular classification, including Cytokeratin+ tumor cells, CD3+CD8+ cytotoxic T cells, CD3+CD4+ helper T cells, CD20+CD3-CD38-B cells, CD38+CD20-Ki67-plasma cells, CD57+CD107a+HLA-DR+ NK cells, CD68+ HLA-DR+ macrophages, Mast Cell Tryptase+ mast cells, Vimentin+SMA+ fibroblasts, CD11b+CD68- granulocytes, Podoplanin+ lymphatic endothelial cells, and CD31+ blood vessel endothelial cells. Spatial distance analysis was conducted using the MACS iQ View Software. The outlines of identified cells were transposed to a Leica Microsystems LMD7 laser microdissection instrument. Cell registration was accomplished by establishing three reference points through the Biology Image Analysis Software (BIAS, Cell Signaling) ^15^. Cells were precisely extracted using the middle pulse function and collected into a 384-well plate. Quadruplicate samples were obtained, each covering a surface area of 37,000 μm².

### MS sample preparation

The collected cells were sedimented at the base of each well through acetonitrile-assisted centrifugation and subsequent vacuum evaporation until completely dried. Subsequently, lysis was performed using 4 μL of 60 mM triethylammonium bicarbonate (TEAB) and 0.01% n-Dodecyl β-D-maltopyranoside (DDM), followed by incubation at 95°C for 60 minutes in a thermal cycler (S1000 with 384-well reaction module, Bio-Rad). To this, 1 µL of 60% acetonitrile was added, and the samples were heated at 75°C for 1 hour. Protein digestion was achieved using 4 ng LysC and 6 ng trypsin overnight at 37°C. The colorectal cancer samples were label-free and directly subjected to LC-MS/MS analysis after protein digestion. The samples were acidified with 1.22 μL of 10% TFA (final concentration 1%), vacuum dried, and stored at -20°C or resuspended in Evosep buffer A (0.1% formic acid) for direct Evotip loading. For samples derived from tonsil cancer tissues, dimethyl labeling was performed following the method described by Thielert and Itang et al ^16^. A parallel bulk tissue slice was digested as the reference channel. The labeling involved sequential addition of 1 μL of 1.5% formaldehyde (CH2O, CD2O, or 13CD2O) and 1 μL of 230 mM sodium cyanoborohydride (NaBH3CN or NaBD3CN), followed by incubation at room temperature for 1 hour on a bench-top mixer. To halt the reaction, 1 μL of 1.43% ammonia was added. The samples were acidified with 1.22 μL of 10% TFA (final concentration 1%), vacuum dried, and stored at -20°C or resuspended in Evosep buffer A (0.1% formic acid) for direct Evotip loading. Evotip loading involved using 25 ng of the bulk tissue as the reference channel (Δ0), combined with the target Δ4 and Δ8 channels ^16^.

### LC-MS

To assess the versatility of our workflow, we applied it to colorectal cancer FFPE sections and analyzed them using the Orbitrap Astral mass spectrometer (Thermo Fisher Scientific) interfaced with the Evosep One LC system (EvoSep). The Whisper40 SPD method was employed, utilizing the Aurora Elite TS analytical column (IonOpticks) and the EASY-Spray™ source. Mobile phases consisted of 0.1% formic acid (FA) in LC–MS-grade water as buffer A and 99.9% acetonitrile/0.1% FA as buffer B. All samples were recorded in DIA mode. The Orbitrap Astral mass spectrometer was utilized for full MS analyses with a resolution setting of 240,000 within a full scan range of 380–980 m/z. The automatic gain control (AGC) for the full MS was adjusted to 500%. The MS/MS isolation window was set to 8 Th, the maximum ion injection time to 14 ms, and the MS/MS scanning range between 380 − 980 m/z. Selected ions were fragmented by higher-energy collisional dissociation at a normalized collision energy (NCE) of 25%. Samples derived from tonsil cancer tissues underwent analysis using the Evosep One LC system (EvoSep) coupled to a timsTOF SCP mass spectrometer (Bruker). The Whisper40 SPD (samples per day) method was applied, employing the Aurora Elite CSI third-generation column with 15 and 75 μm ID (AUR3-15075C18-CSI, IonOpticks) at 50°C within the nanoelectrospray ion source (Captive spray source, Bruker). Mobile phases consisted of 0.1% formic acid (FA) in LC–MS-grade water as buffer A and 99.9% acetonitrile/0.1% FA as buffer B. The timsTOF SCP was operated with an optimized dia-PASEF method, encompassing 8 dia-PASEF scans with variable width and 2 ion mobility windows per dia-PASEF scan. This spanned an m/z range from 300 to 1200 and an ion mobility range from 0.7 to 1.3 V/cm²/s. The mass spectrometer functioned in high sensitivity mode, with an accumulation and ramp time set at 100 ms, a capillary voltage of 1400 V, and collision energy following a linear ramp from 20 eV at 1/K0 = 0.6 V/cm²/s to 59 eV at 1/K0 = 1.6 V/cm²/s.

### MS data analysis

For label-free colorectal samples, the raw files were initially converted to the mzML file format using MSConvert software (Proteowizard) ^40^, with default parameters and ‘Peak Picking’ selected as the filter. Subsequently, mzML files were quantified in DIA-NN v1.8.1 against a human FASTA reference file (2023, UP000005640_9606), employing a direct-DIA approach^41^. DIA-NN search parameters included: Enzyme specificity set to ‘Trypsin/P’ with a maximum of one missed cleavage. Parameters for post-translational modifications like N-terminal methionine excision, methionine oxidation, and N-terminal acetylation were activated, allowing for a maximum of two variable modifications. Precursor FDR was limited to 1%, and both mass and MS1 accuracy were set to 15 p.p.m. Protein inference was conducted using ‘Genes’, with the neural network classifier in ‘Single-pass mode’ and quantification strategy as ‘Robust LC (high precision)’. Cross-run normalization was set as ‘RT-dependent’, while library generation involved ‘IDs, RT and IM Profiling’. Mass accuracy and MS1 accuracy were automatically inferred from the first run, with additional settings such as ‘Use isotopologues’, ‘Match between runs (MBR)’, ‘Heuristic protein inference’, and ‘No shared spectra’ enabled for optimal results.For dimethyl-labeled tonsil cancer samples, the spectral library was constructed using five dda-PASEF single shots from a 50 ng bulk reference peptide. Spectra were searched using FragPipe v18.0 and MSFragger v3.5, Philosopher v4.4.0, and EasyPQP v0.1.32 against a human FASTA reference file (2023, UP000005640_9606) from the UniProt database 19. The DIA_SpecLib_Quant workflow was applied with standard settings, except for fixed modifications where N-terminal and lysine mass shifts of 28.0313 Da were set, and methionine oxidation was defined as a variable modification. Carbamidomethylation was unselected due to samples not undergoing reduction and alkylation. One missed cleavage was permitted. The precursor charge ranged from 2 to 4, the peptide mass range was set from 300 to 1,800, and peptide length ranged from 7 to 30. The resulting library file was utilized in DIA-NN v1.8.1 to analyze the raw files from diaPASEF measurements ^25^. DIA-NN search parameters mirrored those used for label-free samples, with similar settings for protein inference, quantification strategy, and additional optimizations. Additional commands were entered into the DIA-NN command line GUI, including (1) {--fixed-mod Dimethyl, 28.0313, nK}, (2) {--channels Dimethyl, 0, nK, 0:0; Dimethyl, 4, nK, 4.0251:4.0251; Dimethyl, 8, nK, 8.0444:8.0444}, (3) {--original-mods}, (4) {--peak-translation}, (5) {--ms1-isotope-quant}, (6) {--report-lib-info}, and (7) {-mass-acc-quant 10.0}. The processed DIA-NN output report table underwent additional analysis using the ‘RefQuant’ package implemented in Python. The RefQuant output was filtered for ‘Lib.PG.Q.Value’ < 0.01, ‘Q.value’ < 0.01 and ‘Channel.Q.Value’ < 0.15 ^16^. Subsequently, the data was consolidated into protein groups using the MaxLFQ algorithm, implemented in the R package ‘iq’ ^42^.

### Statistical analysis

The proteome datasets underwent filtering to include samples with at least 70% valid values, excluding proteins with more than 30% missing values for subsequent statistical analyses. Remaining missing values were imputed using the K-Nearest Neighbors (KNN) imputation method ^43^. All samples’ intensity values underwent a logarithmic transformation (log2) and were used for principal component analysis (PCA) and identification of differentially expressed proteins. Protein visualization employed volcano plots and heat maps, analyzed using Python and R. For multi-sample comparisons (ANOVA) or pairwise proteomic analyses (two-sided unpaired *t-test*), a protein was deemed significantly differentially abundant if the FDR-adjusted P-value was less than 0.05. Ontology enrichment analysis on the liver proteomics dataset was conducted using ClueGo (v2.5.10), a Cytoscape (v.3.1.1) plug-in, with default settings. A customized reference set comprising around 5000 unique genes (**Figure 3A-B; Figure S3C**) quantified in this study was employed in Fisher’s exact test. Term significance was corrected using the Benjamini–Hochberg method with an FDR threshold of less than 1%. The analysis encompassed the activation of both Gene Ontology term fusion and grouping.

## Acknowledgements

We express our gratitude to Dr. Franziska Neumann, Dr. Lea Bornemann and Dr. Olga Hartwig for their insightful discussions on the MACSima platform. Dr. Yasuko Antoku maintained the MACSima platform at optimal performance. Dr. Lise Mette Rahbek Gjerdrum provided the colorectal cancer tissue samples. Special appreciation goes to Ericka Corazon Itang, Dr. Constantin Ammar, and Dr. Juanjuan Wang for their valuable insights and discussions on multiplex-DIA (mDIA) mass spectrometry analysis. Their expertise greatly contributed to the successful implementation of the mDIA workflow in our study. Discussions with Dr. Thierry Nordmann and Dr. Lise Mette Rahbek Gjerdrum on pathology were immensely constructive. This work was supported by the Novo Nordisk Foundation (grant agreements NNF14CC0001 and NNF15CC0001).

## Supplementary figures

**Figure S1.**
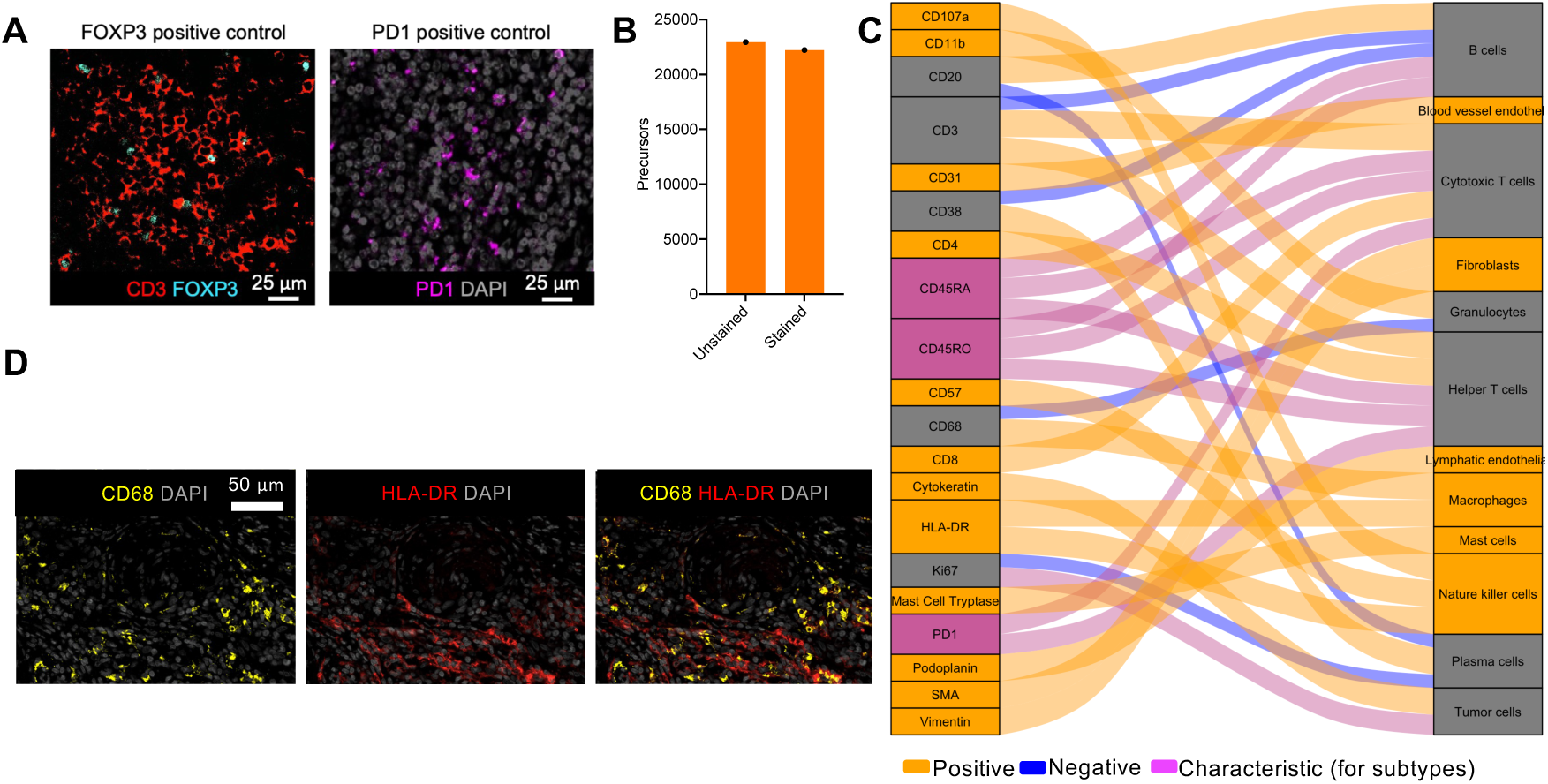
(A) Positive control of FOXP3 and PD1 staining on tonsil tissue, validating the specificity and reliability of the multiplex immunofluorescence staining protocol. (B) Precursor analysis of unstained and 22-plex stained tonsil cancer tissue. (C) Alluvial plot illustrating the markers used in cellular classification, providing an overview of the cell type identification strategy. (D) Representative image depicting CD68 and HLA-DR expression in colorectal cancer tissue.

**Figure S2.**
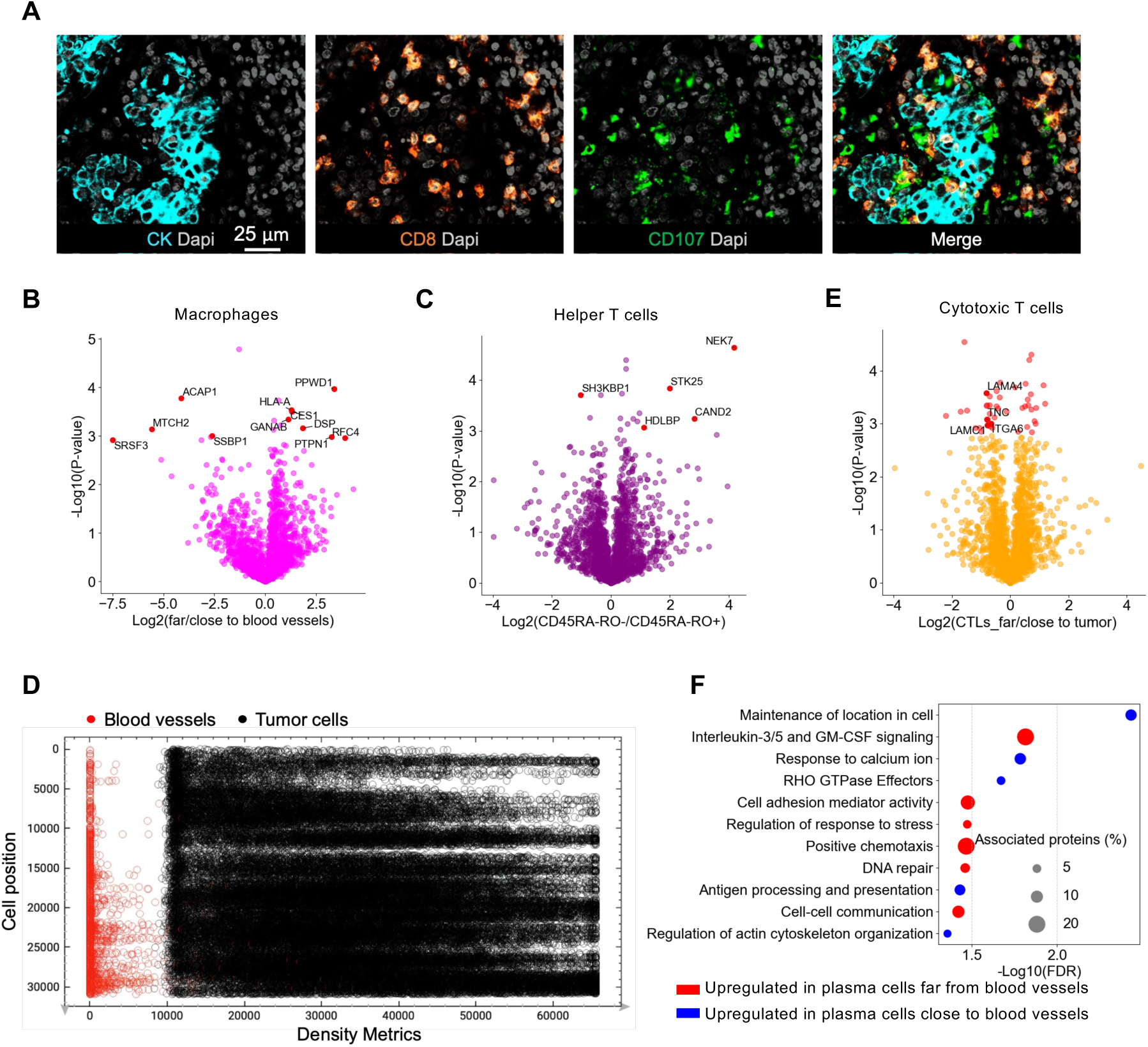
(A) Representative image depicting CD107 expression in tonsil cancer. (B) Volcano plot showing proteomic comparisons between macrophages in regions proximal to and distal from blood vessels. (C) Volcano plot showing proteomic comparisons between CD45RA-CD45RO- (RA-RO-) and RA-RO+ helper T cells (TH) from tonsil cancer. (D) Density metrics displaying blood vessel and tumor cell densities relative to nuclear X position, with higher values indicating greater density. The plot reveals the spatial distribution of blood vessels and tumor cells within the tonsil TME. (E) Volcano plot illustrating proteomic differences in CD45RA-CD45RO- (RA-RO-) cytotoxic T cells (CTLs) based on their proximity to the tumor site in tonsil cancer. (F) Gene Ontology (GO) term analysis of significantly upregulated proteins in plasma cells located far from or close to blood vessels in tonsil cancer.

**Figure S3.**
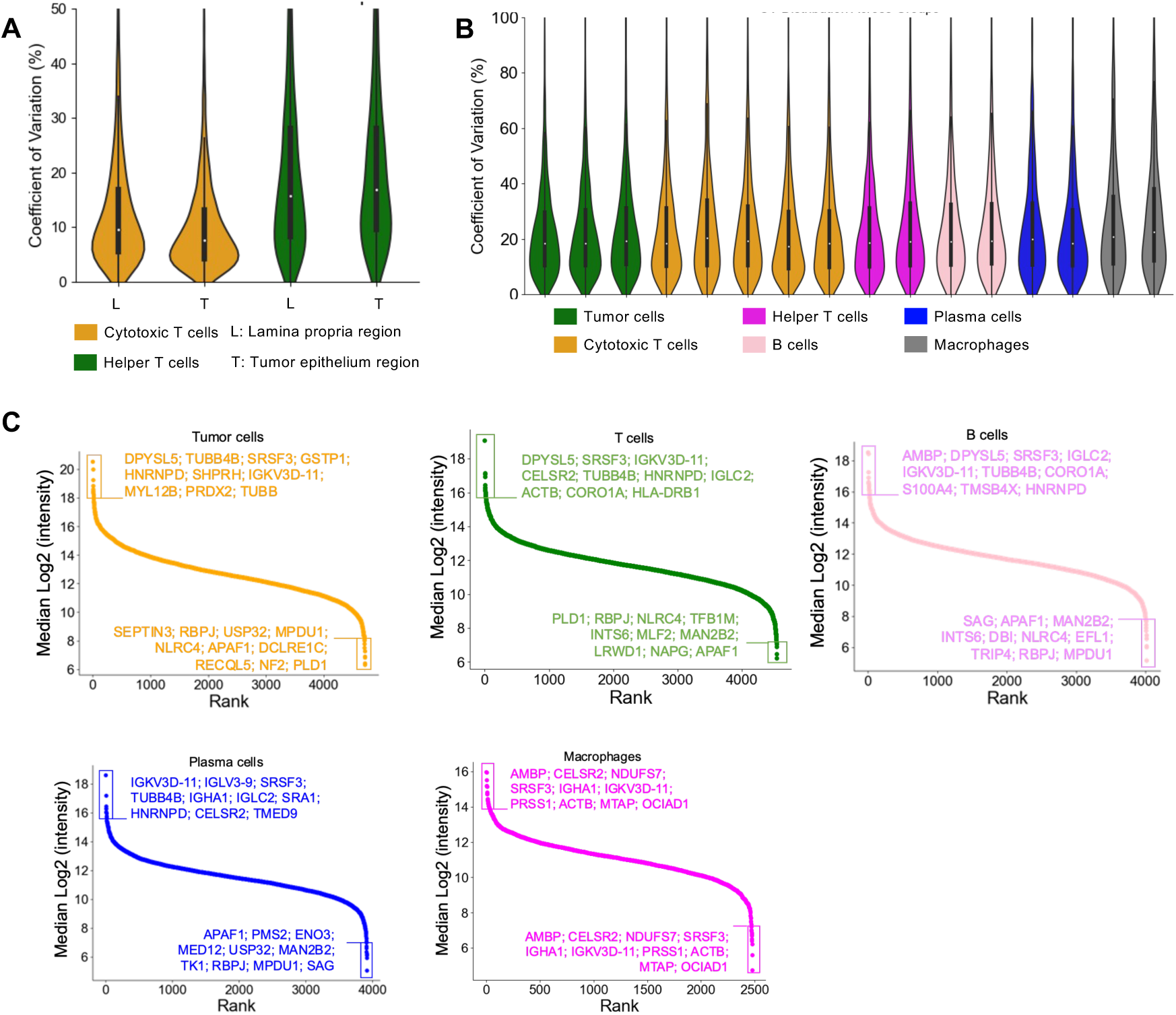
(A-B) Coefficients of variation (%) across specified cell types under respective conditions, demonstrating the reproducibility and consistency of the proteomic data across different cellular subpopulations and experimental conditions. (C) Protein ranking plot of proteins identified in tumor cells, T cells, B cells, plasma cells, and macrophages from tonsil cancer. Proteins are ranked by median transformed intensity values detected by mass spectrometry.

